# VICR: A Novel Software for Unbiased Video and Image Analysis in Scientific Research

**DOI:** 10.1101/2024.06.26.600811

**Authors:** Kyle Kihn, Clementine A.D. Thomas, Tinatin I. Brelidze

## Abstract

In scientific research, objectivity and unbiased data analysis is crucial for the validity and reproducibility of outcomes. This is particularly important for studies involving video or image categorization. A common approach of decreasing the bias is delegating data analysis to researchers unfamiliar with the experimental settings. However, this requires additional personnel and is prone to cognitive biases. Here we describe the Video & Image Cutter & Randomizer (VICR) software (https://github.com/kkihnphd/VICR), designed for unbiased analysis by segmenting and then randomizing videos or still images. VICR allows a single researcher to conduct and analyze studies in a blinded manner, eliminating the bias in analysis and streamlining the research process. We describe the features of the VICR software and demonstrate its capabilities using zebrafish behavior studies. To our knowledge, VICR is the only software for the randomization of video and image segments capable of eliminating bias in data analysis in a variety of research fields.

## Introduction

Maintaining objectivity and minimizing biases are cornerstone principles for ensuring the validity and reproducibility of research study findings. This is especially critical in studies involving the categorization of videos or images, where a prior knowledge of experimental parameters or subjective interpretations can significantly influence the analysis and conclusions. A pervasive challenge with implementing an unbiased analysis is the absence of specialized tools adept at systematically addressing and mitigating the issue of biases [1-3]. Traditional data analysis methods rely on human researchers, who, despite their expertise, are inherently susceptible to cognitive biases [4-8]. These biases can manifest in various forms, such as confirmation bias, where researchers may unconsciously seek information that confirms their preconceptions; hindsight bias, which can lead researchers to view events as more predictable after they have occurred; and anchoring bias, where initial information disproportionately influences subsequent judgments [9-11]. Furthermore, a trained researcher analyzing the experiment, even if not directly involved in its execution, may possess intuitive knowledge about the experimental setup, expected results, and test conditions based on the interactions with other researchers in the same group or field of studies. This indirect knowledge can influence their categorization decisions, compromising the unbiased data analysis and conclusions of the study.

To address these challenges, we developed a novel approach using the Video & Image Cutter & Randomizer (VICR) software (https://github.com/kkihnphd/VICR), which is designed to segment videos and still images according to the user selection and then randomize the sequence of the segments for unbiased analysis. Once the user completes data analysis within the individual segments, the software assigns the analysis results to the corresponding segments and generates the analysis summary. This user-friendly tool allows a single researcher to set up, conduct and analyze studies in a truly blinded manner, eliminating the influence of prior knowledge and expectations on the data categorization process. The benefits of having one researcher conduct the entire study are manifold. It streamlines the research process, reducing the potential for communication errors between different researchers, and ensures a consistent approach to data collection and analysis. Additionally, this reduces the number of researchers needed to perform the study, helping to free up recourses and time in the lab. The software’s capability to randomize video and still image segments ensures that the analysis is conducted without any predetermined order or pattern, thus minimizing the risk of researchers’ biases. Additionally, multiple videos/images can be processed at once allowing for a wide range of test conditions to be randomized simultaneously and analyzed together.

We applied the VICR software to analyze zebrafish behavior in an unbiased manner. Zebrafish have emerged as a valuable model organism for studying various diseases due to their genetic similarity to humans, transparency during early development, and rapid life cycle [4;12-14]. Similar to studies using other animal models, zebrafish behavior studies frequently involve examination of multiple animals under a variety of conditions. For instance, to study epilepsy using zebrafish, seizures are typically induced with different concentrations of pentylenetetrazole (PTZ) [4;13]. For behavior analysis, it is essential to keep the researcher analyzing the data blinded to the PTZ concentration used for each animal. To accomplish this, we uploaded the video recording of zebrafish placed into wells with different concentrations of PTZ into the VICR software, assigned individual wells as different segments and let the software randomize the segments. PTZ-induced seizures were then analyzed for each individual segment that was randomly presented for the analysis. The comparison of the results of the blinded (randomized) data analysis with the results of the unblinded (unrandomized) data analysis by a researcher familiar with the experimental setup revealed significant differences, highlighting the utility of the software in decreasing the biases. Therefore, we expect VICR to be a useful tool for eliminating biases in zebrafish and other animal research, as well as in non-animal studies where randomizing video and still image segments would increase objectivity and reproducibility of conclusions.

## Results

### Overview of VICR Workflow

VICR (Video & Image Cutter & Randomizer) software was developed to facilitate unbiased analysis of video and image data with the workflow illustrated in Fig. 1. In the beginning of the data analysis with the VICR software, the users input videos or still images to be analyzed and select the media type by choosing between video or image inputs (Fig. 1a). VICR accommodates a spectrum of video and image data formats for analysis, including the commonly used MPEG-4 Part 14 (.mp4), Audio Video Interleave (.avi) formats for video analyses, and Portable Network Graphics (.png), Joint Photographic Experts Group (.jpg/.jpeg), and Bitmap (.bmp) for image analyses. The software permits batch selection of data from multiple files of videos or images, seamlessly combining them for blinded analysis. In addition, the user can indicate the desired number of parameters to be analyzed and customize their names, tailoring the software output to the specifics of their study.

**Figure 1:**
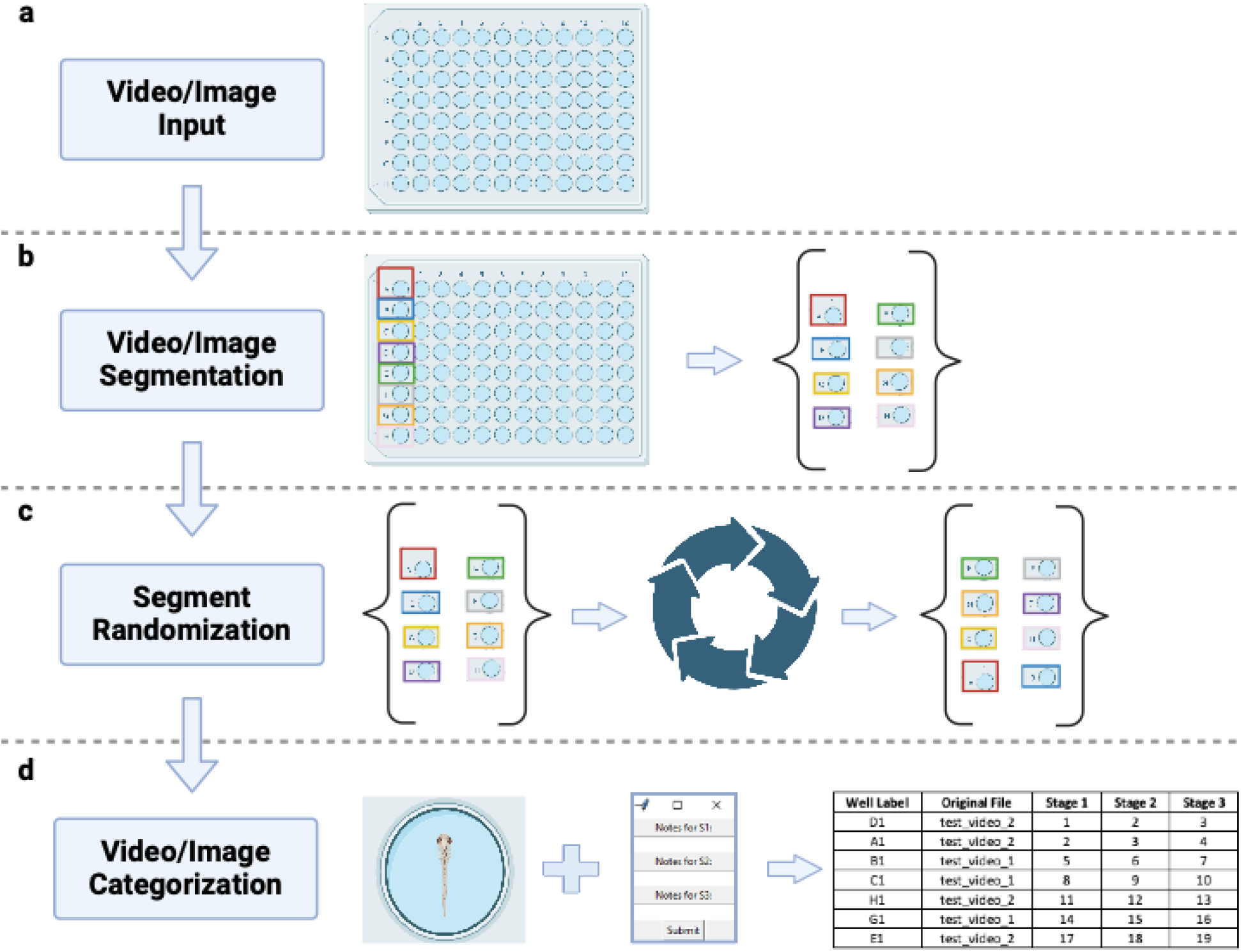
The VICR software workflow described in four principal stages. (a) Media Input: The user initiates the process by importing either video or image files. (b) Media segmentation: The user selects and labels the individual segments within each video or image. (c) Segment Randomization: The user selected segments undergo a randomization process, ensuring unbiased segment sequence presentation for subsequent analysis. (d) Media categorization: In the final step, media segments, randomly presented to the user, are reviewed and categorized by the user. In this stage, the software internally tracks the analysis details blinded to the user. At the end of this stage, the analysis results are saved in a table that can be used for subsequent analysis.

After the media selection and input, the interface guides the user to the segmentation phase (Fig. 1b). Here, the user defines segments of interest by encompassing the target areas in images or initial frames of videos with a rectangle controlled by a computer mouse motion, and labels each segment with a distinct identifier. At present the source media can only be subdivided into rectangular segments. Although more flexibility for outlining the area of interest might be beneficial for some studies, the rectangular rather than flexible border selection helps to mitigate the user bias in the segment selection. Once all the segments are identified, VICR dissects the media into the defined segments, followed by the segment storage and randomization (Fig. 1c). The segments, each bearing the label conferred by the user, are deposited within a results folder, where their sequence is shuffled using the random.shuffle() function from Python’s intrinsic random library (Fig. 1c). The preservation of the randomization sequence during the randomization function operation offers users the flexibility to halt and restart their analysis without compromising the original random order derived from the mixing of all media segments. The results folder can be accessed post-analysis and the individual segments can be re-viewed, if needed.

In the final phase of the workflow, the segments, devoid of any identifying labels, appear for analysis in the software interface in the sequence based on the randomization enacted in the previous step (Fig. 1d). After evaluating each individual segment, the user inputs their observational notes into a dialogue box for each of the parameters (stages) to be analyzed. Although the segment’s label remains concealed from the user during analysis to preclude bias, VICR internally tracks this information. The results of the analysis are saved as a table in an Excel file, which contains the user-defined labels, media source files, and analysis notes for each randomized segment. The described workflow ensures unbiased and systematic data analysis.

### User-Friendly Interface and Operations

The interface of VICR is designed to seamlessly guide the user through the steps of the workflow. The main software window includes buttons for the media type selection and selecting one or multiple media files for analysis (Fig. 2a). Once the selection is completed a follow-up dialog window allows the user to select between new analysis or resuming an old analysis. In addition to the media selection, the user can specify the name of the folder where the selected data segments will be stored, and the number and name of parameters or stages that will be analyzed for each segment (Fig. 2b).

**Figure 2:**
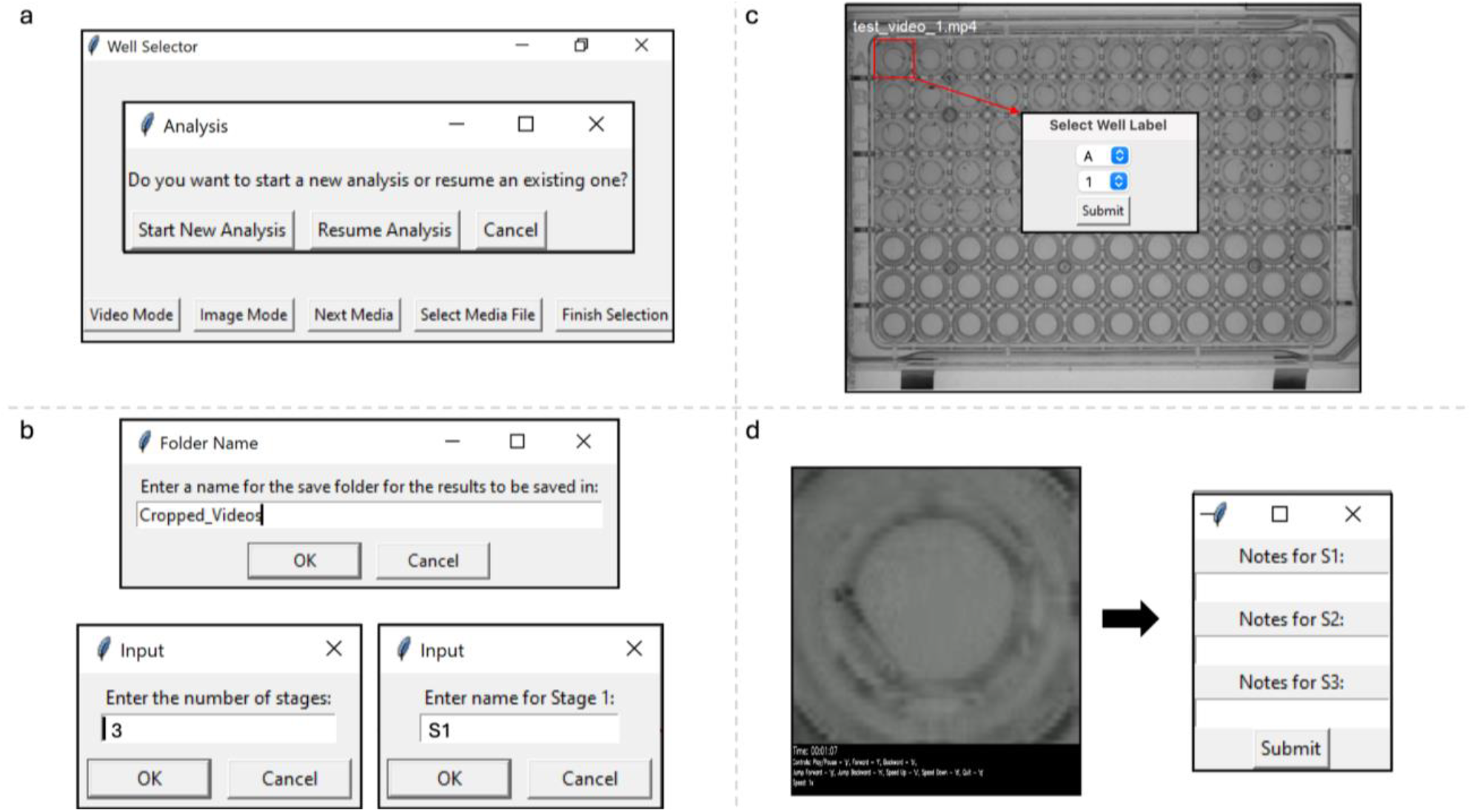
The VICR user interface illustrating the operational workflow. (a) The initial dashboard provides options for starting a new analysis or resuming a previous one. (b) Follow-up dialog windows provide the user with the opportunity to name the folder where the analysis results will be stored, to indicate the number of stages to be analyzed and to name each of the stages. (c) A snapshot of a video frame displaying a 96-well plate where the user can select a specific well; an inset dialog box captures the labeling process for the selected segment. (d, left panel) A user defined segment, that appears in a random order in the software interface, with the video control options listed in the black panel at the bottom of the video. (d, right panel) A dialog box that enables the user to ender analysis information for each of the stages.

VICR’s selection tool enables rectangular delineation of regions within the media files. This is illustrated in Fig. 2c that shows an image of a 96-well plate (first frame of the video) with a zebrafish in each well in rows A-E. The red box demarcates the segment selection that will be used for subsequent examination. To ensure unique identification, the interface prompts the user to assign labels for each of the selected segment using letters (A-Z) and numbers (1-36) (Fig. 2c, insert). The software prevents duplicate labeling, further aiding in accuracy and reliability of the segment selection. However, once selected the naming of the segment cannot be changed for the ongoing analysis. Once the selection of the individual segments of the media has been completed, the software proceeds to the randomization phase, wherein the order of the selected segments is shuffled using Python’s intrinsic random library, guaranteeing an absence of any discernible sequence. This step is meticulously logged, not just to support an unbiased analysis, but also to maintain the sequence of randomization of the segments, should the analysis be paused and resumed later. This is especially important and convenient when large data sets and/or multiple media files are analyzed together as such an analysis could be time consuming and require multiple sessions. The randomization process is typically quick for still images. However, for long video recordings, the segmentation and randomization can take substantial time depending on the user’s computer specifications.

Once the randomization process is over, the segments are displayed in the order of randomization for analysis (Fig. 2d, left). The software does not affect the resolution of the media used for the segmentation. Therefore, the resolution of the displayed segments will be the same as the resolution of the initial media at the same magnification. Several options are available to facilitate the analysis of video segments, including playback speed adjustments, fast forwarding, rewinding, and play/pause options. The options are summarized on the black panel at the bottom of the video recording with the video time prominently displayed at the top of the panel. After the user is finished with the media evaluation, the notes or conclusions of the analysis for the user-defined parameters/stages can be entered into a dialog box (Fig. 2d, right).

At the end of the analysis of all selected segments, the software inputs the analysis results into an Excel spreadsheet that consolidates the user’s observations, including notes and numerical evaluations, with their associated segment labels and source media files. The user can then use the results of the unbiased analysis for making subsequent conclusions for the study based on the knowledge of specific experimental conditions. This approach should increase the rigor of scientific conclusions by enabling unbiased data analysis for experiments based on image or video assessments.

### Application in Zebrafish Behavior Studies

To demonstrate VICR’s suitability for unbiased analysis, we conducted zebrafish behavior studies involving the induction of seizures with varying concentrations of PTZ, a GABA_A_ receptor inhibitor used to induce seizure-like activity in zebrafish and rodents [4;15]. In zebrafish, PTZ-induced seizure-like behavior is characterized by three distinct stages. Stage I is associated with a sudden increase in swim activity, stage II with a rapid circular movement around the well edge and stage III with a whole-body convulsion followed by a loss of posture and a period of immobility, as depicted in Fig. 3a [4;5;16]. To observe the PTZ-induced seizure-like behavior, 7 days post fertilization (dpf) zebrafish larvae were placed in a 96-well plate with a mesh insert (one fish per well) and monitored for a period of 2 minutes in fish water (acclimation). To ensure a rapid and even application of PTZ, the mesh insert with the fish was then transferred into a new 96-well plate with wells containing only fish water (control) or containing fish water with 2.5 mM, 5 mM or 15 mM PTZ. Following the transfer, the fish were monitored for a period of 20 minutes to observe PTZ-induced behavior. To determine the latency to the three stages, the video recording of the fish behavior was analyzed using the VICR software, involving the segmentation of the whole plate recording and randomization of the segments (blinded analysis) and by directly analyzing the whole plate video recording without any measures to blind the experimental conditions used (unblinded analysis). Both blinded and unblinded analysis was performed by the same researcher who conducted the behavioral study and was familiar with the experimental conditions. The results of the blinded and unblinded analysis for the three stages are compared in Fig. 3. Overall, the determined latencies to the three stages were consistent with previous reports [4], however, a close examination revealed notable differences between blinded and unblinded analysis.

**Figure 3:**
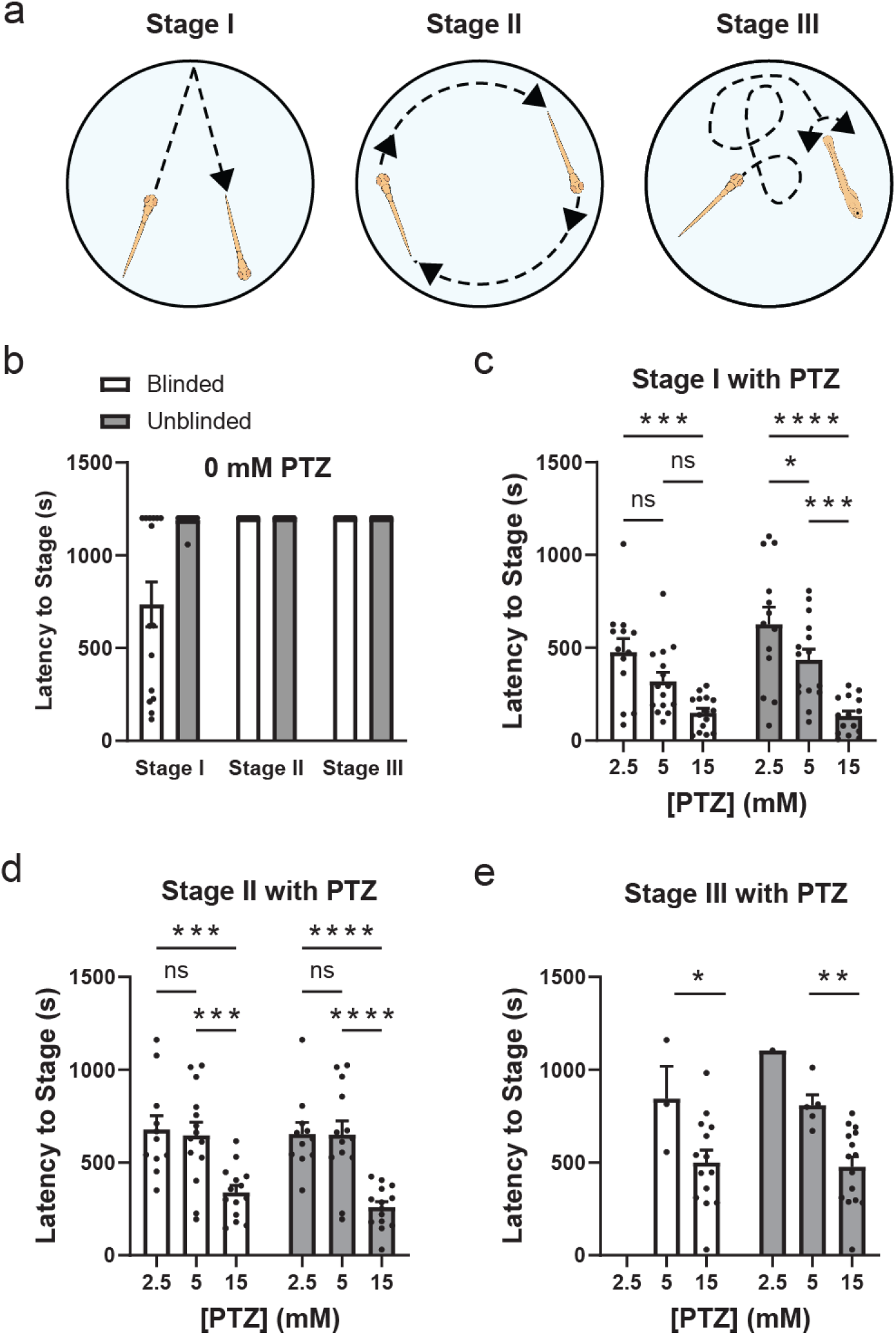
Comparative analysis of latency periods for zebrafish seizure stages under varying concentrations of PTZ. (a) Schematic representation of the three seizure-associated behavior stages in zebrafish. (b) Latency times to reach Stages I, II and III in the absence of PTZ based on the blinded (white bar graphs) and unblinded (grey bar graphs) analysis. (c-e) Latency times to reach Stages I, II and III of seizure-associated behavior, respectively, with the indicated PTZ concentrations based on the blinded (white bar graphs) and unblinded (grey bar graphs) analysis. Statistical annotations indicate levels of significance between blinded and unblinded analysis within each PTZ group, with ‘ns’ denoting not significant, and asterisks * indicating *P*-values of <0.05 (*), <0.01 (**), <0.001 (***), and <0.0001 (****).

The comparison between blinded and unblinded analysis for Stage I revealed a difference in the absence of PTZ (the control conditions) (Fig. 3b). For Stage I, based on the unblinded analysis no fish reached a seizure-like behavior during the 20 minutes of the experiment, while based on the blinded analysis 60% of fish reached Stage I seizures. For stages II and III, there was no difference between the blinded and unblinded analysis results in the absence of PTZ and no fish displayed Stage II or Stage III seizures during the 20 minutes of the experiment. For all three stages blinded and unblinded analysis showed a tendency for a concentration dependent decrease in the latency to each stage with the increase in the PTZ concentration. However, overall for all stages the statistical significance in the decrease in the seizure latency with the increase in the PTZ concentration was more significant based on unblinded than blinded analysis. For instance, for the latency to Stage I seizures, there was no statistically significant difference for blinded analysis between 2.5 mM and 5 mM PTZ, and also between 5 mM and 15 mM PTZ, with statistically significant difference observed only between 2.5 mM and 15 mM PTZ (Fig. 3c). In contrast, for unblinded analysis statistical significance in the decrease in latency to Stage I was observed with each increase in the PTZ concentration. For Stage II analysis, the statistical significance was higher for the latency to seizures between 5 and 15 mM PTZ, and 2.5 and 15 mM PTZ, for unblinded than blinded analysis (*P* < 0.001 versus *P* < 0.0001 according to the ANOVA test) (Fig. 3d). Similarly, for Stage III analysis the statistical significance was higher for the latency to seizures between 5 and 15 mM PTZ for unblinded than blinded analysis (*P* < 0.05 versus *P* < 0.01 according to the ANOVA test) (Fig. 3e). Majority of fish did not reach Stage III seizures with 2.5 mM PTZ and, therefore, this concentration was excluded from the statistical analysis. Taken together, the comparison of statistical significance for blinded and unblinded analysis shows that unblinded analysis would result in statistically more significant conclusions. This illustrates an inherent bias in the analysis of seizure stages in zebrafish and highlights the ability of VICR software to minimize the bias.

## Discussion

Here we describe the VICR software designed to eliminate bias in analyzing data presented in video or still images. To our knowledge, this is the first software with these capabilities. The software allows segmentation of video and still images by the user. The segments are then randomized and presented to the user for analysis devoid of any user-defined and identifying information. This ensures an unbiased analysis of the experiments contained in each segment even by the user familiar with the experimental conditions. The video recordings and still images that could be analyzed with VICR do not require any additional modification and the software can combine analysis of multiple media files of the same type in one analysis session. The user friendly design of VICR and the simple workflow should further facilitate its application for unbiased data analysis for a variety of research, including animal behavior and microscopy.

We have demonstrated the applicability of the VICR software to zebrafish behavior studies. We examined the PTZ-induced seizure associated behavior in zebrafish. For these experiments PTZ was added at different concentrations and latency to the three seizure stages was determined using blinded analysis with VICR and compared to the results of unblinded analysis performed by the same researcher. For the data analysis with VICR, the video recordings were segmented into recordings of individual wells, the segments were randomized, and analyzed in a blinded manner (Fig. 1). It is expected that the latency to seizure stages would decrease with the increase in the PTZ concentration. Therefore, the bias in this analysis could be introduced by misinterpreting fish movement based on the expected effect of PTZ on the seizures. This possibility was further confirmed by the comparison of our results of the blinded and unblinded analysis. For both blinded and unblinded analysis, the determined latency to the three seizure stages was overall consistent with previous reports [4]. However, comparison between blinded and unblinded analysis results revealed a statistically significant difference favoring the expected results for the unblinded analysis. Not surprisingly, the difference between the blinded and unblinded results was the most statistically significant for Stage I (Fig. 3b and 3c), which is arguably the most subjective of the seizure stages to score as it involves sudden movement somewhat similar to what is observed in PTZ-free conditions. Although less prominent, similar bias towards decreased latency was observed for Stages II and III in the presence of PTZ for unblinded relative to the blinded analysis (Fig. 3d and 3e). Overall, the comparison of the blinded and unblinded results indicates that there are inherent biases potentially introduced through prior knowledge of experimental conditions, influencing the interpretation of results for behavior studies. The blinded data analysis with the VICR software allows for an accurate and objective assessment of seizure stages, resulting in the minimization of bias which is vital for upholding the credibility of research findings.

In addition to eliminating bias in zebrafish behavior analysis, VICR simplifies the overall analysis process by keeping track of the inputs in the excel sheet. VICR also allows researchers to exit and return to the analysis at any point, which eliminates possible confusion in analysis. By focusing solely on the fish present in the individual segments being analyzed, the VICR software eliminates the distractions that may arise from the movement of fish in other wells on the plate.

The analysis of behavioral experiments with VICR illustrated with the zebrafish model in our study, should be applicable to behavioral studies of other model organisms monitored in delineated regions under different conditions. In addition to unbiased analysis of behavioral research, the VICR software can also be used for other fields involving visual image analysis. For instance, in environmental research VICR can help with the impartial classification of imagery from natural habitats and wildlife. In oncology, VICR can facilitate unbiased analysis of cell differentiation, viability, migration and other characteristics in response to a drug treatment in situations when researchers have to manually evaluate experimental outcomes. For these studies, frequently conducted on cell cultures in multi-well plates, VICR should ensure the unbiased assessment of the therapeutic potential of antitumorigenic drugs.

Taken together, VICR should increase the objectivity and reproducibility of the research conclusions in a variety of fields involving video/image analysis of multiple objects under different conditions. The capability of the software to blind the analysis enables a single researcher familiar with the research conditions to oversee the entire research cycle, from experimental setup to data analysis, removing bias in data analysis and conserving laboratory resources.

## Methods

### Software Design and Implementation

The Video & Image Cutter & Randomizer (VICR) software was developed as a Python-based application, utilizing the Tkinter library for the graphical user interface (GUI) and random.shuffle() function from Python’s intrinsic random library. VICR’s workflow is divided into four main functionalities: media input, segmentation, randomization, and categorization. The media input functionality allows input of the media (still image or video) and pausing or resuming the analysis. The GUI facilitates multi-file uploads and categorization into respective media types. The segmentation functionality allows users to subdivide the source media into segments by outlining the area of interest with a rectangle with the cursor on first frame of a video file or the source image, and labeling of the outlined segments with unique identifiers. The segment randomization functionality utilizes random.shuffle() function from Python’s built-in random library to randomize the order of the media segments. The randomization protocol ensures that the sequence of segment analysis is devoid of any user-induced patterns, thus enforcing an unbiased study structure. The software records the randomization order to enable continuity in cases where analysis is paused and resumed at a later time. The categorization functionality allows the user to observe the selected segments, which appear in a randomized order devoid of any identifying labels, and enter the conclusions into a dialog window. The software accepts a variety of video formats, including MPEG-4 Part 14 (.mp4) and Audio Video Interleave (.avi), as well as image formats such as Portable Network Graphics (.png), Joint Photographic Experts Group (.jpg/.jpeg), and Bitmap (.bmp).

Upon completion of the analysis, VICR compiles the results into an Excel spreadsheet. The output file systematically lists the user-defined labels for each segment, the original file names, and the associated notes from the categorization process. This enables a straightforward transition to further data analysis based on the knowledge of specific experimental conditions.

The detailed step-by-step instructions on how to install and use the software, and descriptions of the major functions of the software, are provided in the accompanied VICR User Guide and VICR User Tutorial. The required components to run VICR include Python 3.x, OpenCV for Python (cv2), Pillow (PIL), Tkinter, NumPy, and OpenPyXL and can be installed following the steps outlined in the accompanied User Guide. These components are needed for video and image processing, GUI operation, and data management functionalities within the software. VICR is compatible with major operating systems (Unix, MacOS, WindowsOS).

### Code Availability

The source code for VICR, as well as the user guide and tutorial, can be found at https://github.com/kkihnphd/VICR.

### Zebrafish husbandry

Zebrafish were maintained at 28°C under a 14:10 h light to dark cycle. All animal procedures were conducted in accordance with NIH guidelines for the care and use of laboratory animals and approved by the Georgetown University Institutional Animal Care and Use Committee.

### Zebrafish behavior

Behavior experiments were conducted on 7dpf zebrafish larvae placed in a clear 96-well plate with a mesh insert (Fisher Scientific, Cat # MANMN4010). Each well had one fish placed in 450 μl of fish water without or with PTZ (Sigma-Aldrich). 60 zebrafish larvae were used in total. The fish water contained 0.3 g/L of sea salt in distilled water. For the experiments the 96-well plate with zebrafish larvae was placed in the DanioVision observation chamber (Noldus, Germany) and zebrafish movement was tracked with the EthoVision XT video tracking software (Noldus, Germany). The fish was first monitored for 2 minutes in PTZ-free fish water (acclimation period) and then the mesh insert with the fish was transferred into a new plate with 15 mM PTZ in columns 1-3, 5 mM PTZ in columns 4-6, 2.5 mM PTZ in columns 7-9 and no PTZ in columns 10-12. With this experimental design, 15 zebrafish larvae were analyzed for each PTZ concentration and the PTZ-free control condition.

The fish behavior was then monitored for an additional 20 minutes. The two video recordings of the behavior experiment, corresponding to the 2-min acclimation and 20-min of the PTZ application periods, are available upon request.

The data for the latency to seizure stages were analyzed using two-way ANOVA followed by a Tukey’s multiple comparisons test.

## Supporting information

VICR user guide

VICR user tutorial

## Acknowledgements

This work was supported by the National Cancer Institute grant R01CA252969 (T. I. B.), and National Institute of General Medical Sciences grants R01GM124020 (T.I.B.) and T32GM142520 (C.A.D.T.). The Animal Model Zebrafish Shared Resource is partially supported by the National Cancer Institute grant P30-CA051008.

## Author contributions

K.K., C.A.D.T. and T.I.B. conceived and designed the study. K.K. developed the code for VICR. K.K., C.A.D.T. and T.I.B. tested the software and analyzed the results. K.K. and T.I.B. wrote the manuscript with the input from all authors.

