## Supplementary material for "VICR: A Novel Software for Unbiased Video and Image Analysis in Scientific Research": VICR user guide

### User Guide for VICR

**Initial Installation**

Installing Anaconda Navigator

1. **Download Anaconda Installer**: Go to the Anaconda website and download the Anaconda installer for your operating system.
2. **Run the Installer**: Double-click the downloaded file to launch the installer. Follow the prompts to complete the installation. Choose the default settings for an easy setup.
3. **Verify Installation**: Open Anaconda Navigator from the Start menu (Windows) or Launchpad (macOS). For Linux, use the command **anaconda-navigator** in the terminal.

Setting Up the Environment

1. **Create a New Environment**: In Anaconda Navigator, go to the "Environments" tab and click "Create". Name your environment (e.g., **vicr_env**) and select Python version 3.8 or later. Click "Create".
2. **Activate the Environment**: Once the environment is created, click on the play button next to it and select "Open Terminal" to activate the environment.

Running the Code

1. **Install Required Packages**: In the activated terminal, install the necessary packages by running **pip install opencv-python pillow numpy pandas openpyxl tk**.
2. **Launch the Script**: Navigate to the directory containing your script (**VICR.V.1.3.py**) by typing **cd** command followed by the directory path (e.g. e.g. cd C:\Users\Username\Downloads\VICR_Tutorial). Run the script by typing **python VICR.V.1.3.py**.

**Running the Code After Initial Installation**

1. **Open Anaconda Navigator**: Launch Anaconda Navigator from the Start menu (Windows) or Launchpad (macOS).
2. **Activate Environment**: Go to the "Environments" tab, select your environment (e.g., **vicr_env**), and click the play button to activate it.
3. **Open Terminal**: Click on the play button next to the environment and select "Open Terminal".
4. **Run the Script**: Navigate to the directory containing your script (**VICR.V.1.3.py**) by typing **cd** command followed by the directory path (e.g. e.g. cd C:\Users\Username\Downloads\VICR_Tutorial). Run the script by typing **python VICR.V.1.3.py**.

**Using the Code**

Features

- **Video and Image Analysis**: The code allows for segmentation of the video and still images, and then randomization of the video and image segments for blinded analysis.
- **Cropping and Labeling**: Users can manually crop and label regions of interest in videos and still images.

How to Use

- **Select Media**: Choose between Video Mode and Image Mode to begin analysis.
- **Manual Cropping**: Use the mouse to draw a rectangle around the region of interest and label it. A new directory based of the names of your input files will be automatically

created in the directory of the loaded files. This will serve as the save directory for the code. Files in this directory should not be altered by the user if they wish to resume the analysis at a later time.

- **Video Controls**: Use the video player controls to play/pause, forward, backward, jump forward or backward, and speed down or up the video.
- **Categorization**: Enter the categorization results/notes (e.g. time) for each stage (parameter) when prompted.
- **Saving Results**: The results are automatically saved to an Excel file in the same directory as the media. The code autosaves after each categorization entry.
- **Resuming Analysis**: If the analysis is interrupted, the code can be resumed by running it again. It will automatically detect the last analyzed file and continue from there.
- **Quitting Mid-Analysis**: To quit the analysis before completion, close the launch terminal. Ensure to close the terminal before closing the displayed image or video window to avoid errors.

**<u>Outputs and Results</u>**

- **Cropped Videos/Images**: A folder that contains cropped videos or images of the segments from the original media, labeled according to the user input.
- **Randomization Order**: A hidden text file (**Randomization_Order.txt**) listing the order in which the media files were analyzed.
- **Analysis Results (Excel)**: An Excel file (**Blinded_Analysis_Results.xlsx**) containing the labels and categorization times/notes for each cropped image or video. Each row corresponds to a cropped segment of the video or image, with columns for the well label, original file name, and categorization times/notes for each stage.
- **Well to Original Mapping**: A hidden text file (**.well_original_mapping.txt**) mapping each well label to the original file path, used for resuming analysis.

By following this guide, users can install, run and effectively utilize the VICR tool for categorizing and analyzing video and image data in a blinded manner. The accompanied User Tutorial provides more detailed step-by-step instructions on the software use.
