## Supplementary material for "VICR: A Novel Software for Unbiased Video and Image Analysis in Scientific Research": VICR user tutorial

### User Tutorial for VICR

**Initial Installation (this section is from the User Guide for VICR and is included here for convenience)**

Installing Anaconda Navigator

1. **Download Anaconda Installer**: Go to the Anaconda website and download the Anaconda installer for your operating system.
2. **Run the Installer**: Double-click the downloaded file to launch the installer. Follow the prompts to complete the installation. Choose the default settings for an easy setup.
3. **Verify Installation**: Open Anaconda Navigator from the Start menu (Windows) or Launchpad (macOS). For Linux, use the command **anaconda-navigator** in the terminal.

Setting Up the Environment

1. **Create a New Environment**: In Anaconda Navigator, go to the "Environments" tab and click "Create". Name your environment (e.g., **vicr_env**) and select Python version 3.8 or later. Click "Create".
2. **Activate the Environment**: Once the environment is created, click on the play button next to it and select "Open Terminal" to activate the environment.

Running the Code

1. **Install Required Packages**: In the activated terminal, install the necessary packages by running **pip install opencv-python pillow numpy pandas openpyxl tk**.
2. **Launch the Script**: Navigate to the directory containing your script (**VICR.V.1.3.py**) by typing **cd** command followed by the directory path (e.g. e.g. cd C:\Users\Username\Downloads\VICR_Tutorial). Run the script by typing **python VICR.V.1.3.py**.

**Running the Code After Initial Installation (this section is from the User Guide for VICR and is included here for convenience)**

1. **Open Anaconda Navigator**: Launch Anaconda Navigator from the Start menu (Windows) or Launchpad (macOS).
2. **Activate Environment**: Go to the "Environments" tab, select your environment (e.g., **vicr_env**), and click the play button to activate it.
3. **Open Terminal**: Click on the play button next to the environment and select "Open Terminal".
4. **Run the Script**: Navigate to the directory containing your script (**VICR.V.1.3.py**) by typing **cd** command followed by the directory path (e.g. e.g. cd C:\Users\Username\Downloads\VICR_Tutorial). Run the script by typing **python VICR.V.1.3.py**.

**Tutorial for Video and Image Categorization and Analysis (VICR) Tool**

Prerequisites:

- Ensure that you have the VICR tool installed and set up as per the user guide.
- Make sure that the test files (**test_image_1.png**, **test_image_2.png**, **test_video_1.mp4**, **test_video_2.mp4**) are present in the provided folder named **VICR_Tutorial**.

**Tutorial for Video Analysis:**

1. **Launch the Tool**:
    - Open Anaconda Navigator and activate your environment (e.g., **vicr_env**).
    - Open a terminal and navigate to the **VICR_Tutorial** folder.
    - Run the script: **python VICR.V.1.3.py**.
    - The **Well Selector** window should appear, as shown below:

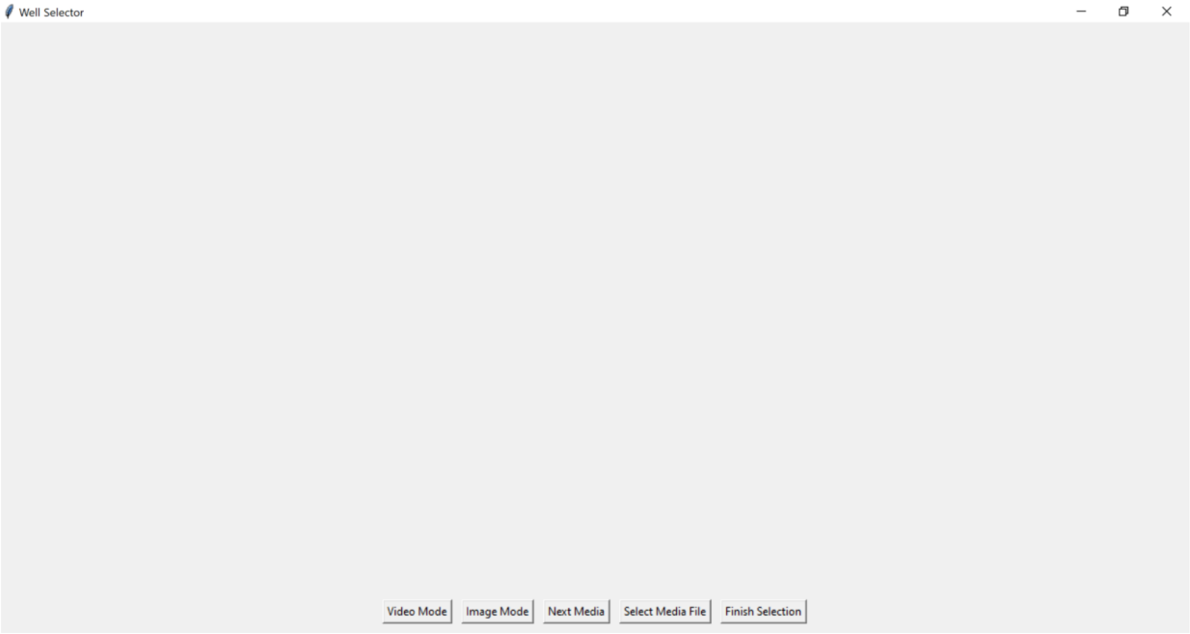


2. **Select Video Mode**:
    - Choose the "**Video Mode**" button in the “**Well Selector**” window.
    - The two left most buttons will change from “**Select Media File**” and “**Finish Selection**” to “**Select Video File**” and “**Finish Video Selection**”.
3. **Select and Categorize Images**:
    - Click the “**Select Video File**” button. This will open a dialogue window to select analysis options: “**Start New Analysis**” to start a new analysis session, “**Resume Analysis**” to continue previously paused analysis session or “**Cancel**” to switch to the image analysis mode, if desired. For the purposes of the tutorial, click “**Start New Analysis**” button. This will open a file navigator window.

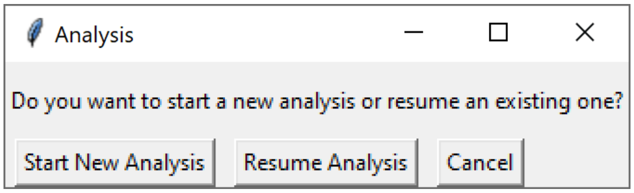

- Navigate to your tutorial folder.  VICR can analyze single or multiple video recordings.  If analyzing only one video recording, select the desired video file and then hit the **Open** button.  For the tutorial we will be analyzing two video recordings provided in the **VICR_Tutorial** folder.  Therefore, select **test_video_1.mp4** and **test_video_2.mp4** files by pressing Cntrl button on the keyboard while clicking on the desired files.  Hit the **Open** button on the file navigator.

  Note: The two video files in the **VICR_Tutorial** folder contain the same video recordings that were given different file names for the purposes of the tutorial.  Also, the video recordings were intentionally short for the purposes of the tutorial.  A longer (20 min) recording corresponding to the actual experiment can be found attached to the main manuscript and can also be used for testing the software.
- A **Folder Name** dialog box will appear to name the folder where the cropped video segments will be saved. We will use name **Cropped-Videos** for the purposes of the tutorial.

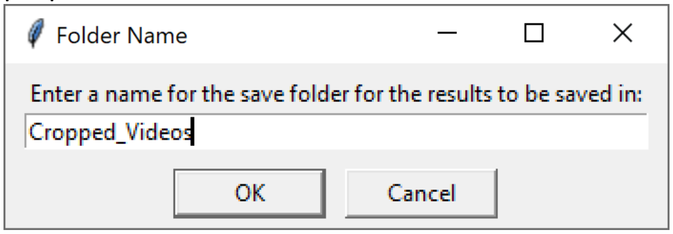


- The dialog box will disappear and an **Input** dialogue box will appear for you to enter the desired number of stages for analysis.  For the tutorial we will be analyzing 3 epilepsy stages in zebrafish.  Therefore, enter 3 into the box and hit "**OK**".

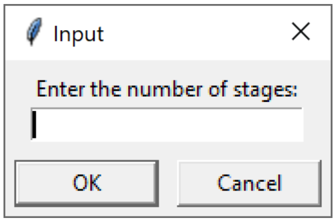


  Note: In general, the number of stages corresponds to the number of experimental parameters that need to be analyzed.
- You will then be prompted to name each of the stages. For this tutorial we will name them S1, S2, and S3.
- Enter S1 into the dialogue box and hit "**OK**", Repeat for S2 and S3.

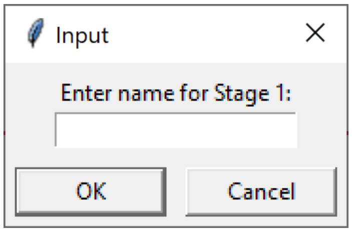


- Once each desired stage has been named, the tool will open **test_video_1.mp4** and display the first frame of the video, as well as the name of the video in the top right corner. Use the mouse to draw a rectangle around a region of interest and label it as prompted.

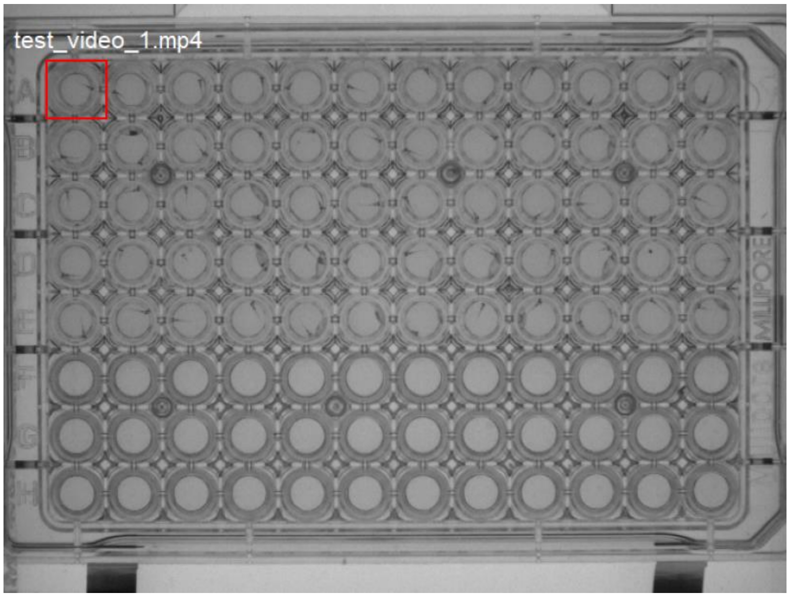


Note 1: Once the selection is finished a dialog box appears to confirm the selection. Press the **Confirm** button. A follow-up dialog box then appears asking if the selection area is correct. If satisfied with the selection, choose **Yes**, or choose **No** and reselect the area. Once the selection is confirmed it cannot be changed for the ongoing analysis.

Note 2: Labels cannot be the same. If you try to label a segment the same as an existing label the code will prompt you to change the current label. Repeat the above step until all desired areas are selected and labeled.

Note 3: We strongly advise to crop all of the desired segments from the beginning to increase the power of randomization.

- To progress to cropping the next video hit the "**Next Media**" button. This will display the first frame of **test_video_2.mp4**.

Note 1: Once you hit this button you cannot go back and change the cropping for the **test_video_1.mp4.**

Note 2: Since the segment (well) labels cannot be the same, the labels for the segments in the second video have to be distinct from the ones in the first video. For instance, we advise to label well A1 in the second video as A13 to make it distinct from the label of the well in the same position in the first video.

- Repeat the cropping and labeling steps for the second video.

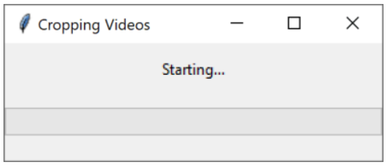


- Once cropping is complete select the "**Finish Video Selection**" button which will begin the video(s) segmentation process. A **Cropping Videos** dialogue window saying **Starting....** will appear and stay visible until the cropping is complete.

Note: For long video recordings, the segmentation and randomization can take substantial time depending on the user's computer pacifications. We advise to avoid using other programs in the background to speed up the process and avoid potential crushing of the computer.

- Once the code had finished cropping according to the user's selection, the cropped segments are randomized and will be randomly displayed one at a time in a new window.

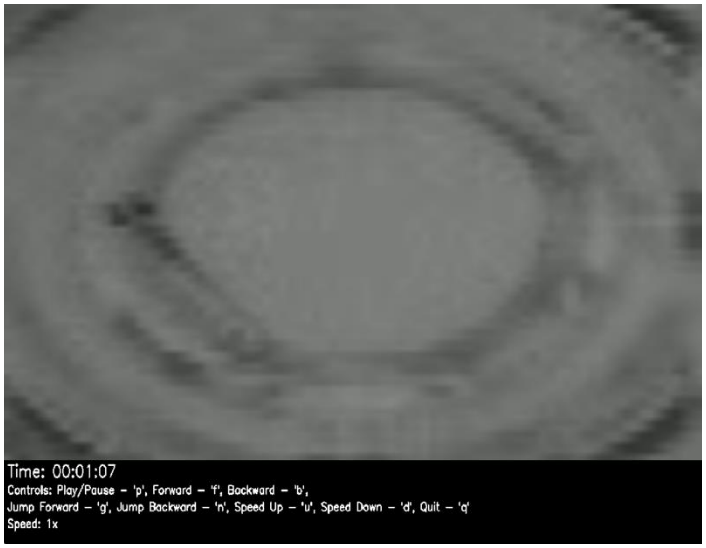


Note: The initially size of the cropped video window most likely will be too large for analysis and should be resized by dragging the corners of the window to the desired size.

- The video for each cropped segment will run in a continuous loop. The user has many options to control the video, which are detailed on the video itself, including pause ("**p**"), forward ("**f**"), backward ("**b**"), jump forward or jump backward ("**g**" and "**n**", correspondingly), speed up ("**u**") or slow down ("**d**"). The forward and backward operations move the video by 24 frames in the corresponding direction, while the jump forward and backward move the video by 1440 frames in the corresponding direction. In a typical 24 frame/sec recording these operations would correspond to 1 second or 1 minute jumps in the forward or backward direction. The speed of the video is also indicated, with **Speed: 1x** corresponding to the real time of the recording. To restart the video after pausing, press pause ("**p**") again. Once the user had determined the stage time or desired notes for each stage hit quit ("**q**") to quit the video. This will close the video and open up a dialogue box to enter the categorization. Fill in the desired information for each stage and then hit the "**Submit**" button.

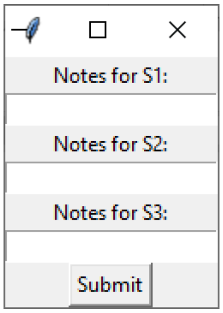


Note: The user may choose to enter just numbers corresponding to the analysis results (e.g. times) and/or more detailed notes.

- This will then have the next video appear for analysis. Repeat the process until all cropped segments are analyzed (at which point the program will automatically close). To exit mid analysis, exit the terminal. To avoid any issues only close the program if a video is displayed, do not exit if the categorization dialog box is active.
- The code auto saves the cropped video segments in the **Cropped_Videos** folder located in **VICR_Tutorial** as defined by the user in the previous steps of the tutorial.

Note: If several videos will be analyzed separately make sure to give unique names to the folders with cropped videos to prevent overwriting.

- The results of the blinded analysis will then be deposited in the **VICR_Tutorial** folder as an excel file (**Blinded_Analysis_Results. xlsx**). The excel spreadsheet will display the values entered by the user for each of the stages for the corresponding segments. An example of the spreadsheet shown below corresponds to the blinded analysis results for the selection of two wells from the first test video and three wells from the second test video, with the corresponding analysis notes for each of the three stages as indicated.

| Well Label | Original File | S1 | S2 | S3 |
|---|---|---|---|---|
| A1 | test_video_1 | 1 | 2 | 3 |
| A15 | test_video_2 | 12 | 2 | 3 |
| A17 | test_video_2 | 1 | 2 | 3 |
| A13 | test_video_2 | 1 | 2 | 3 |
| A1 | test_video_1 | 1 | 2 | 3 |

Note: Once the blinded results are generated, the corresponding cropped segments can be open individually and viewed from the **Cropped_Videos** folder and reanalyzed, if desired.

4. **Exit and Resume Analysis**:
   - To pause the analysis before completing categorization of all of the cropped segments, close the launch terminal to exit the analysis.

     Note: Analysis of at least one cropped segment has to be completed before pausing the analysis session. Otherwise, the selection of the cropped segments will be lost.
   - Once ready to resume the analysis, relaunch the tool from the terminal: **python VICR.V.1.3.py**.
   - Choose the "**Video Mode**" button when the software launches and the "**Well Selector**" window opens.
   - The two left most buttons will change from "**Select Media File**" and "**Finish Selection**" to "**Select Video File**" and "**Finish Video Selection**".
   - Click the "**Select Video File**" button. This will open a dialogue window. Click "**Resume Analysis**" to resume the analysis.
   - A navigation window will open. Select **Cropped-Videos** folder created from the paused session in the **VICR_Tutorial** folder and hit open. This will resume the analysis by pulling up the next video segment in its randomization order.
   - Continue the analysis for each cropped video segment, as described above.
   - Once all cropped segments are analyzed the program will automatically close and the results of the blinded analysis will be deposited in the **VICR_Tutorial** folder as an excel file (**Blinded_Analysis_Results. xlsx**).

     Note: Do not modify files in the **VICR_Tutorial** folder if the analysis is paused as the code uses these files to resume analysis.

**Tutorial for Image Analysis:**

1. **Launch the Tool**:
   - Open Anaconda Navigator and activate your environment (e.g., **vicr_env**).

- Open a terminal and navigate to the **VICR_Tutorial l** folder.
- Run the script: **python VICR.V.1.3.py**.
- The **Well Selector** window should appear, as shown below:

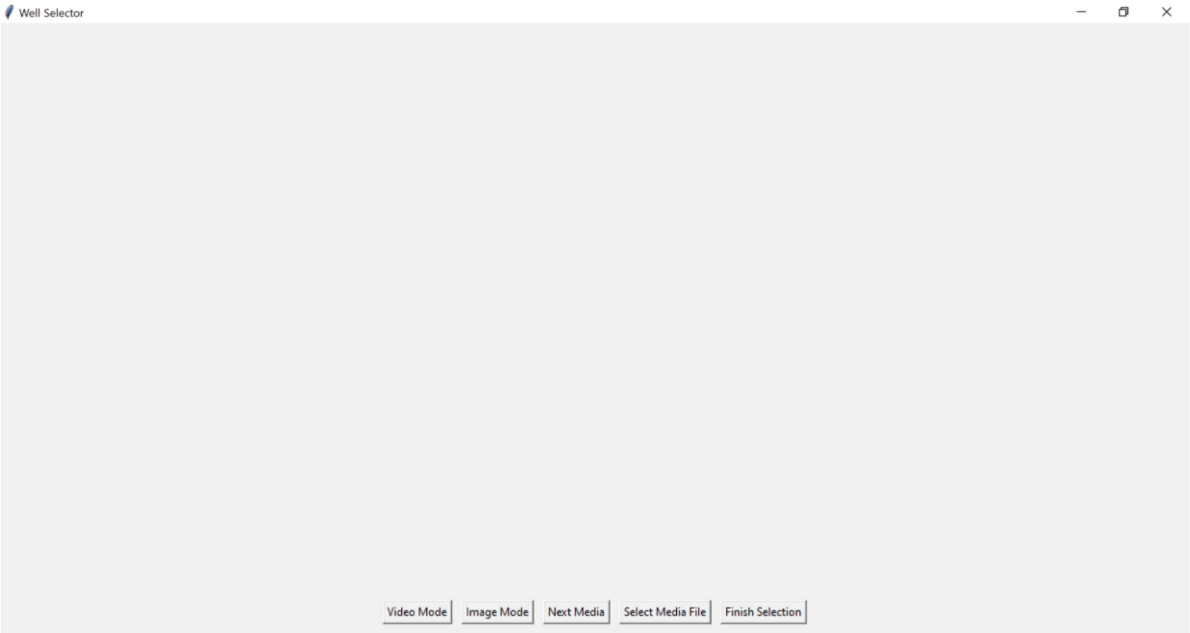


2. **Select Image Mode**:
   - Choose the "**Image Mode**" button in the “**Well Selector**” window.
   - The two left most buttons will change from “**Select Media File**” and “**Finish Selection**” to “**Select Image File**” and “**Finish Image Selection**”.
3. **Select and Categorize Images**:
   - Click the “**Select Image File**” button. This will open a dialogue window to select analysis options: “**Start New Analysis**” to start a new analysis session, “**Resume Analysis**” to continue previously paused analysis session or “**Cancel**” to switch to the video analysis mode, if desired. For the purposes of the tutorial, click “**Start New Analysis**” button. This will open a file navigator window.

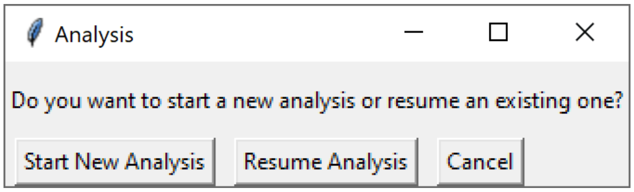


   - VICR can analyze single or multiple image files. If analyzing only one image file, select the desired image file and then hit the **Open** button. For the tutorial we will be analyzing two image files provided in the **VICR_Tutorial** folder. Therefore, select **test_image_1.png** and **test_video_2.png** files by pressing Cntrl button on the keyboard while clicking on the desired files. Hit the **Open** button on the file navigator.

     Note 1: The image files in the **VICR_Tutorial** folder have a “.png” extension, which is the default format for the images. Therefore, to display image files with a different extension select either the appropriate extension or **All files** in the drop-down window.

Note 2: The two image files in the **VICR_Tutorial** folder contain the same images that were given different file names for the purposes of the tutorial.

- A **Folder Name** dialog box will appear to name the folder where the cropped image segments will be saved. We will use **Cropped-Images** for the purposes of the tutorial.

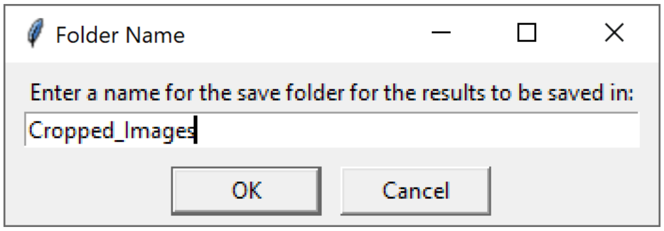


- The file navigator window will disappear and an **Input** dialogue box will appear for you to enter the desired number of stages (parameters) for analysis. For the tutorial we will be analyzing 3 stages. Therefore, enter 3 into the box and hit “**OK**”.

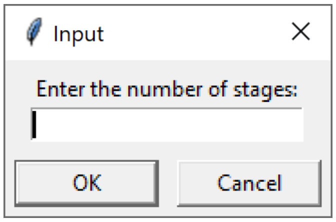


Note: In general, the number of stages corresponds to the number of experimental parameters that need to be analyzed.

- You will then be prompted to name each of the stages. For this tutorial we will name them S1, S2, and S3.

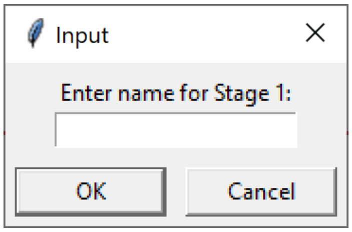


- Once each desired stage has been named, the tool will open **test_image_1.png** and display the image, as well as the name of the image in the top right corner. Use the mouse to draw a rectangle around a region of interest and label it as prompted.

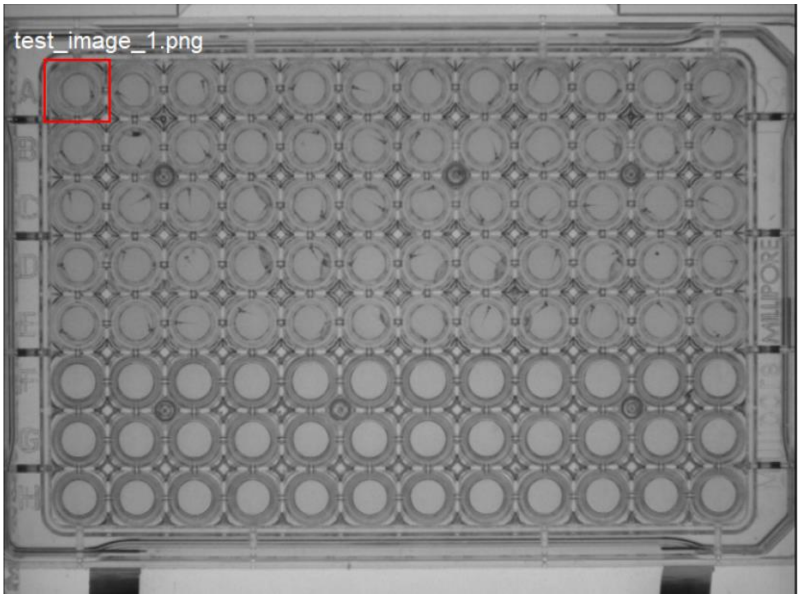


Note 1: Once the selection is finished a dialog box appears to confirm the selection. Press the **Confirm** button. A follow-up dialog box then appears

asking if the selection area is correct. If satisfied with the selection, choose **Yes**, or choose **No** and reselect the area.

Note 2: Labels cannot be the same. If you try to label a segment the same as an existing label the code will prompt you to change the current label.

- Repeat the above step until all desired areas are selected and labeled.

  Note: We strongly advise to crop all of the desired segments from the beginning to increase the power of randomization.

- To progress to cropping the next image hit the "**Next Media**" button. This will display **test_image_2.png** image.

  Note 1 : Once you hit this button you cannot go back and change the cropping for the **test_image_1.png.**

  Note 2: Since the segment (well) labels cannot be the same, the labels for the segments in the second image have to be distinct from the ones in the first image. For instance, we advise to label well A1 in the second image as A13 to make it distinct from the label of the well in the same position in the first image.

- Repeat the cropping and labeling steps for the second image.
- Once cropping is complete select the "**Finish Image Selection**" button which will begin the image(s) segmentation process. A dialogue window saying **Starting....** will appear and stay visible until the cropping is complete.

  Note: Image cropping is typically very fast and the appearance of the dialogue window might be very brief to notice for most of the image cropping tasks, unless large number of images are analyzed.

- Once the code had finished cropping according to the user's selection, the cropped segments are randomized and will be randomly displayed one at a time in a new window.

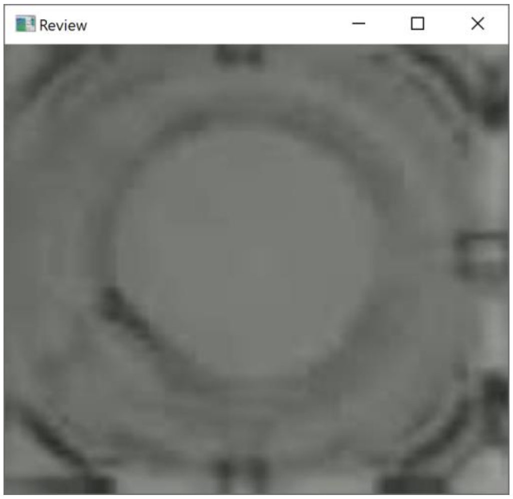


Note: The initially size of the cropped image window most likely will be too large for analysis and should be resized by dragging the corners of the window to the desired size.

- Once the user had determined the desired categorization notes for each stage hit quit ("**q**") to quit the image. This will close the image and open up a dialogue box to enter the categorization. Fill in the desired information for each stage and then hit the "**Submit**" button.

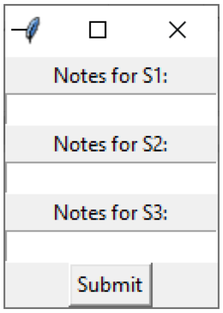


  Note: The user may choose to enter just numbers corresponding to the analysis results and/or more detailed notes.

- Once the categorization notes for one cropped segment are entered then the next randomized cropped segment will appear for categorization.  Repeat the process until all segments are categorized (at which point the program will exit).
- To exit mid analysis, exit the terminal. To avoid any issues only close the program if an image is displayed.  Do not exit if the categorization dialog box is active.
- The code auto saves the cropped video segments in the **Cropped_Images** folder located in **VICR_Tutorial** as defined by the user in the previous steps of the tutorial.

  Note: If several images will be analyzed separately make sure to give unique names to the folders with cropped Images to prevent overwriting.
- The results of the blinded analysis will then be deposited in the **VICR_Tutorial** folder as an excel file (**Blinded_Analysis_Results. xlsx**).  The excel spreadsheet will display the values entered by the user for each of the stages for the corresponding segments.  An example of the spreadsheet shown below corresponds to the blinded analysis results for the selection of two wells from the first test image and three wells from the second test image, with the corresponding analysis notes for each of the three stages as indicated.

| Well Label | Original File | S1 | S2 | S3 |
|---|---|---|---|---|
| A1 | test_video_1 | 1 | 2 | 3 |
| A15 | test_video_2 | 12 | 2 | 3 |
| A17 | test_video_2 | 1 | 2 | 3 |
| A13 | test_video_2 | 1 | 2 | 3 |
| A1 | test_video_1 | 1 | 2 | 3 |

  Note: Once the blinded results are generated, the corresponding cropped segments can be open individually and viewed from the **Cropped_Images** folder and reanalyzed, if desired.

4. **Exit and Resume Analysis**:
   - To pause the analysis before completing categorization of all of the cropped segments, close the launch terminal to exit the analysis.
     Note: Analysis of at least one cropped segment has to be completed before pausing the analysis session.  Otherwise, the selection of the cropped segments will be lost.
   - Once ready to resume the analysis, relaunch the tool from the terminal: **python VICR.V.1.3.py**.
   - Choose the "**Image Mode**" button when the software launches and the “**Well Selector**” window opens.
   - The two left most buttons will change from “**Select Media Fil**e” and “**Finish Selection**” to “**Select Image File**” and “**Finish Image Selection**”.
   - Click the “**Select Image File**” button. This will open a dialogue window. Click “**Resume Analysis**” to resume the analysis.
   - A navigation window will open.  Select **Cropped-Images** folder created from the paused session in the **VICR_Tutorial** folder and hit open. This will resume the analysis by pulling up the next image segment in its randomization order.
   - Continue the analysis for each cropped image segment, as described above.

- Once all cropped segments are analyzed the program will automatically close and the results of the blinded analysis will be deposited in the **VICR_Tutorial** folder as an excel file (**Blinded_Analysis_Results. xlsx**).

  Note: Do not modify files in the **VICR_Tutorial** folder if the analysis is paused as the code uses these files to resume analysis.
